## Supplementary figures and images for "Contribution of behavioural variability to representational drift"

### Figure 1--figure supplement 1

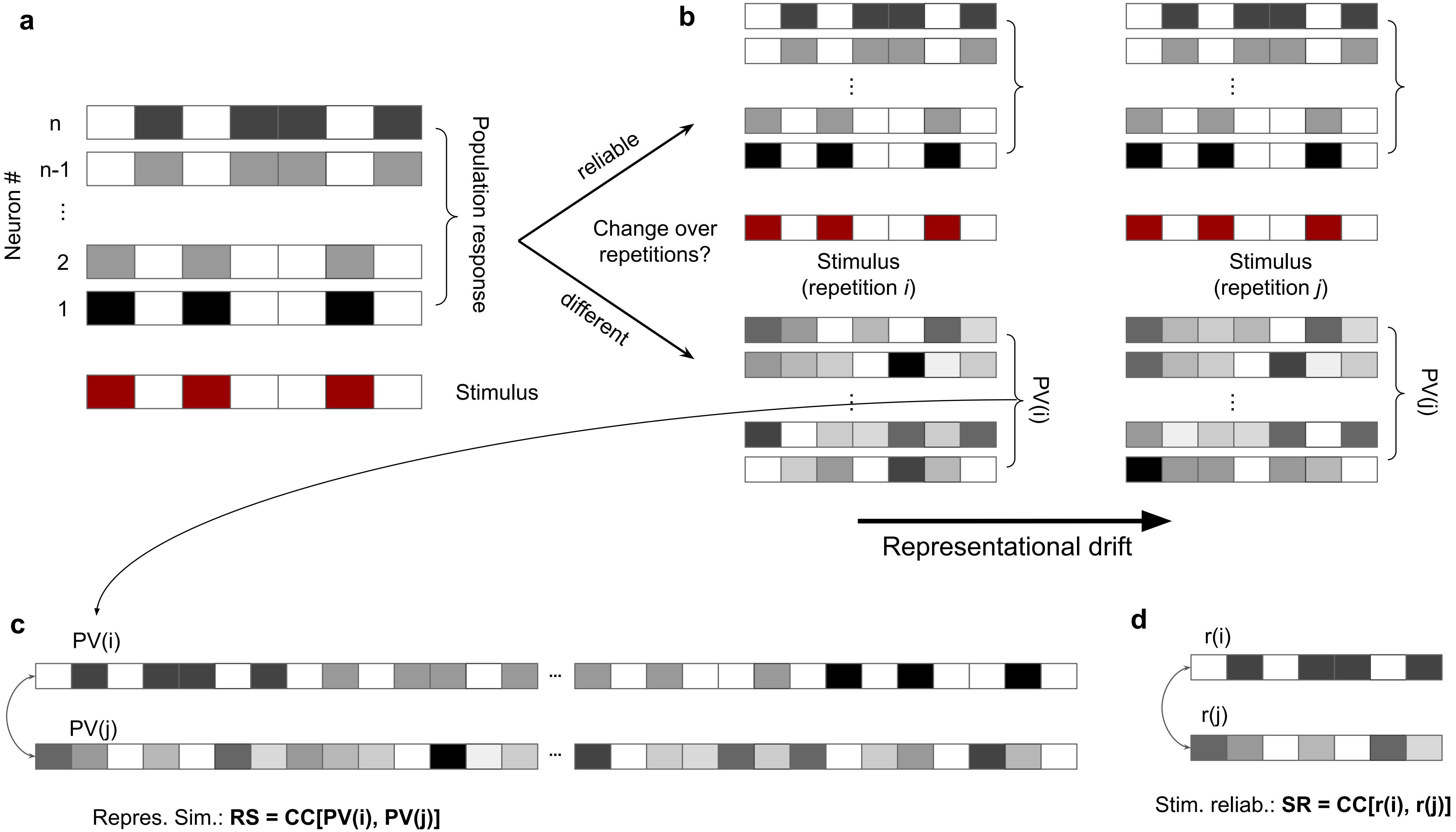

### Figure 1--figure supplement 2

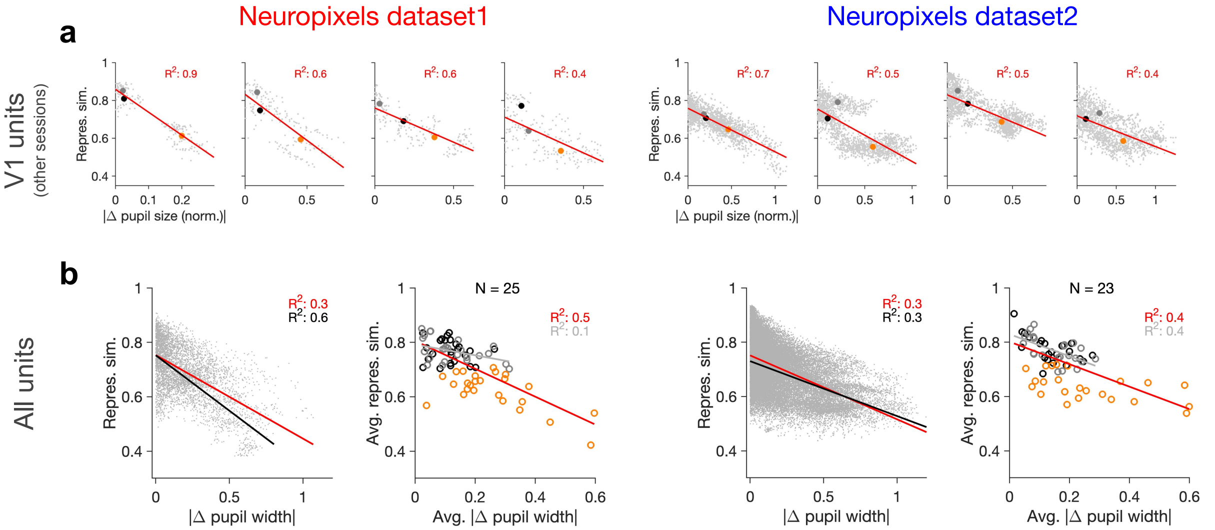

### Figure 1--figure supplement 3

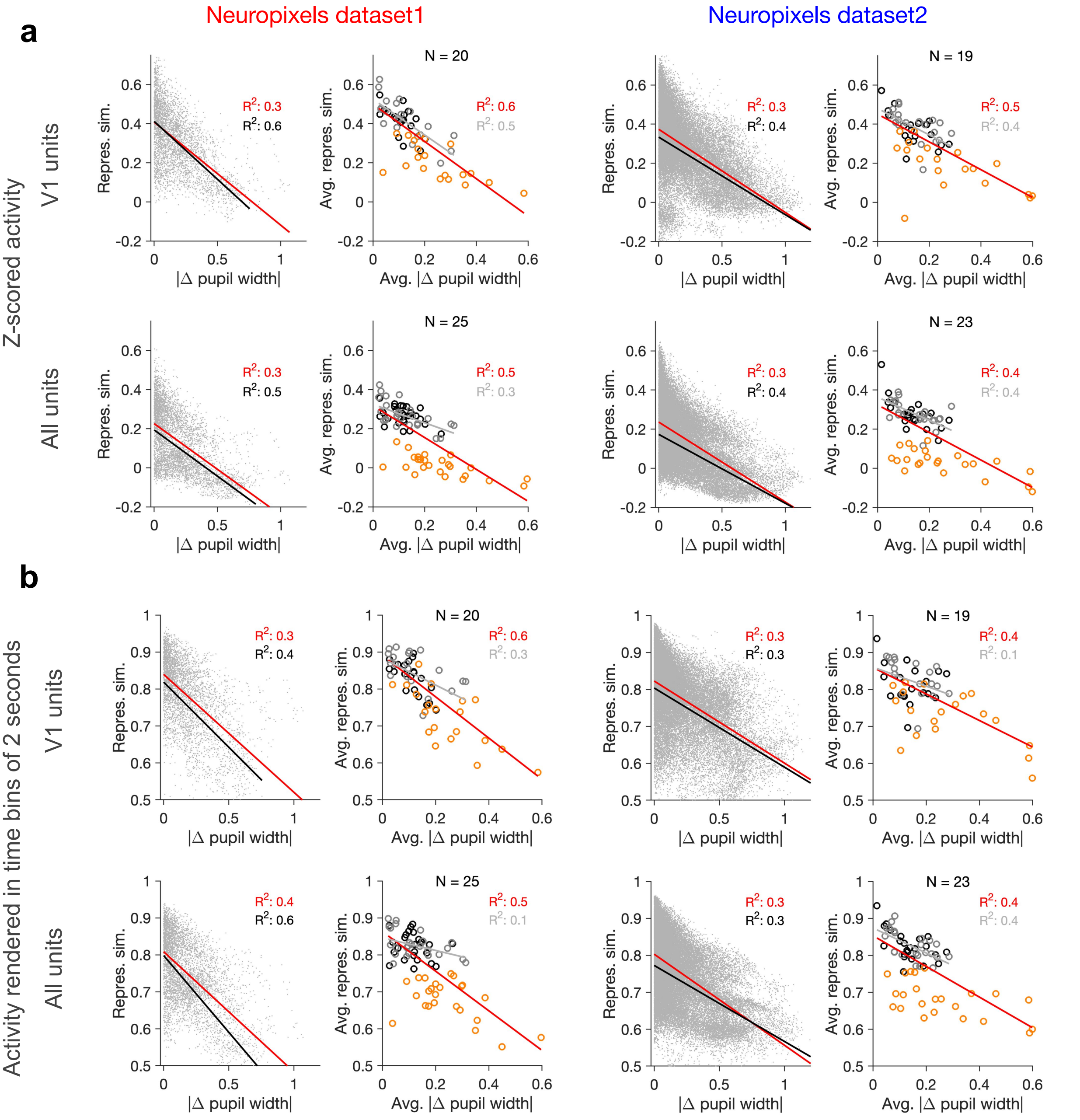

### Figure 1--figure supplement 4

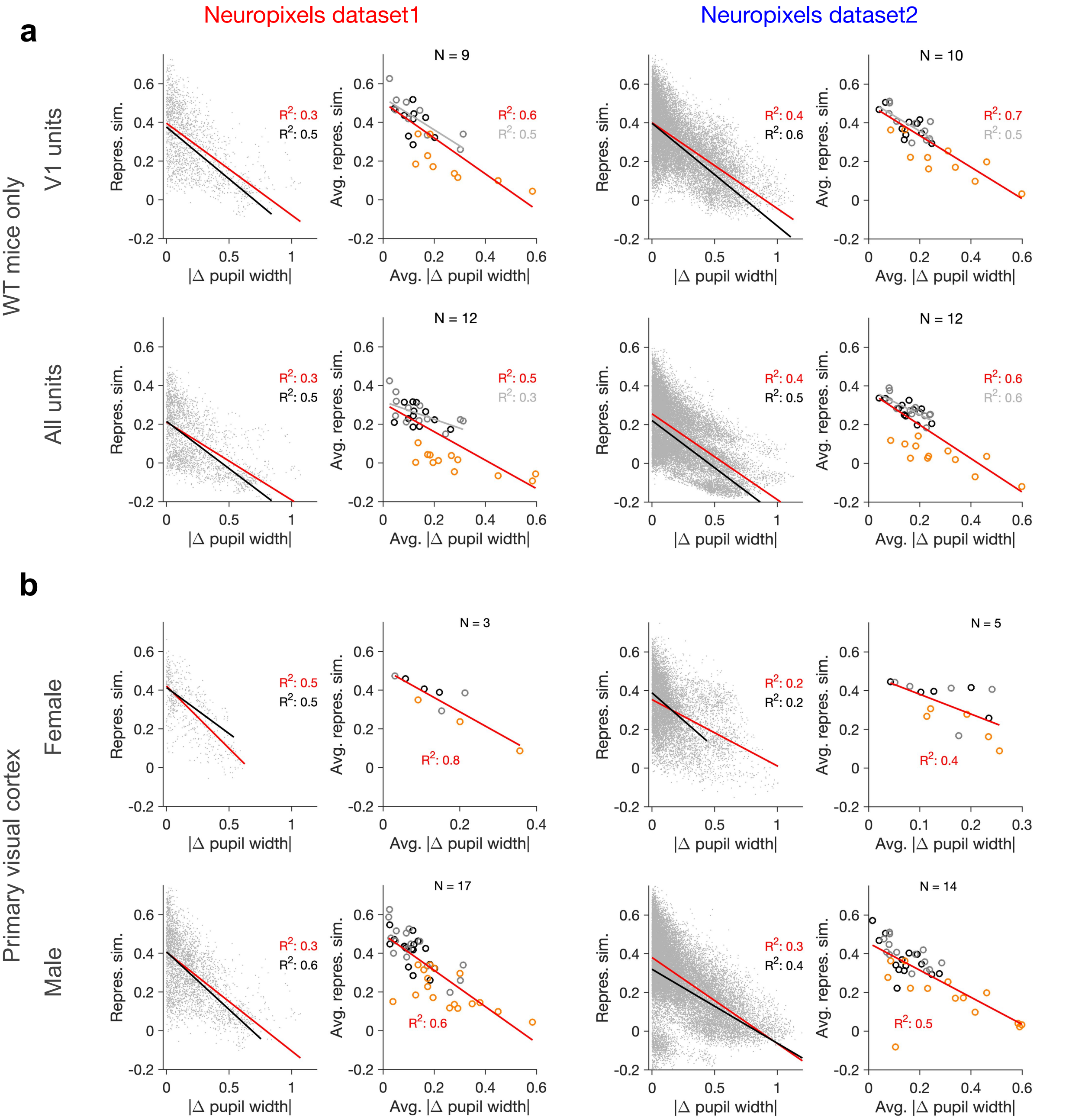

### Figure 1--figure supplement 5

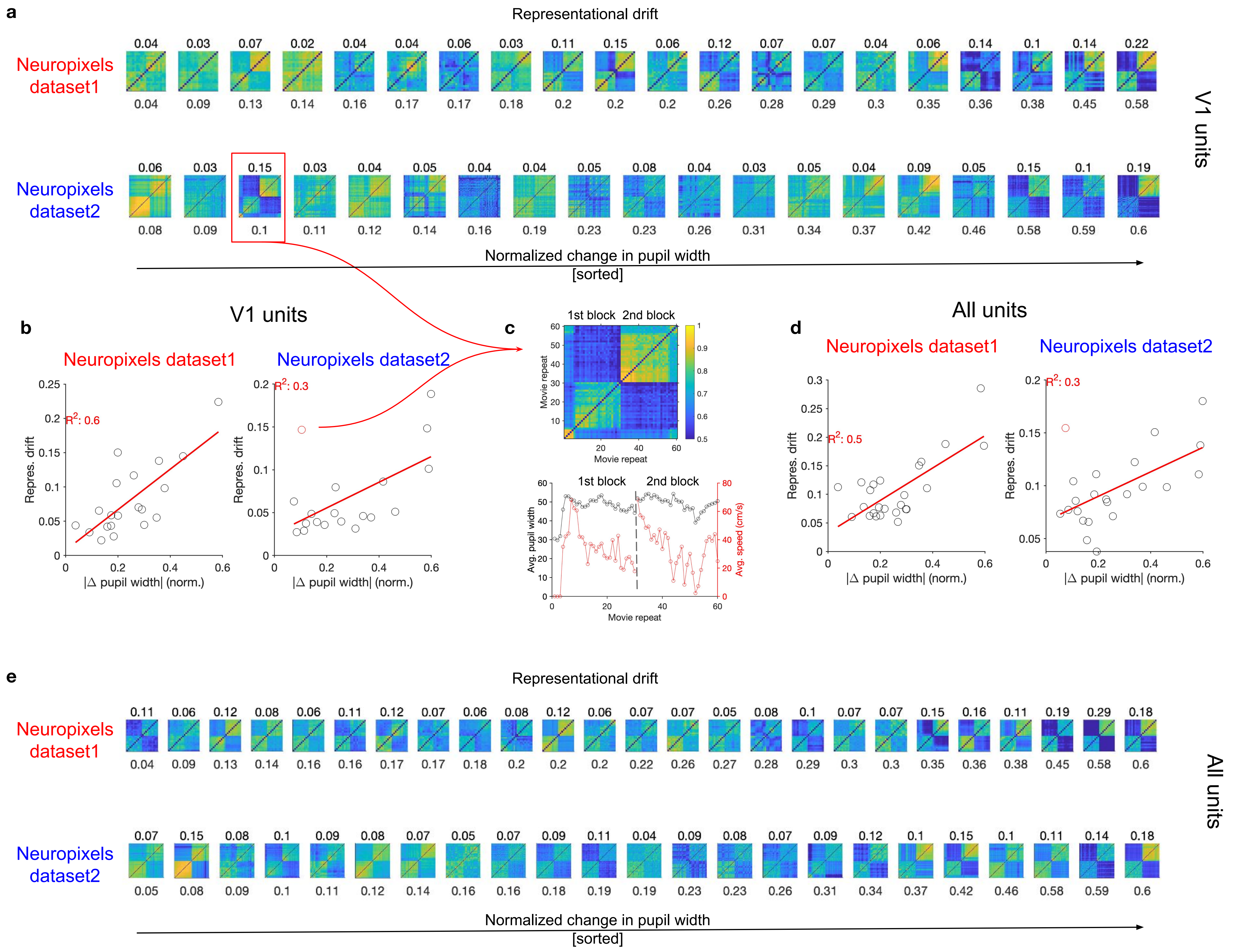

### Figure 2--figure supplement 1

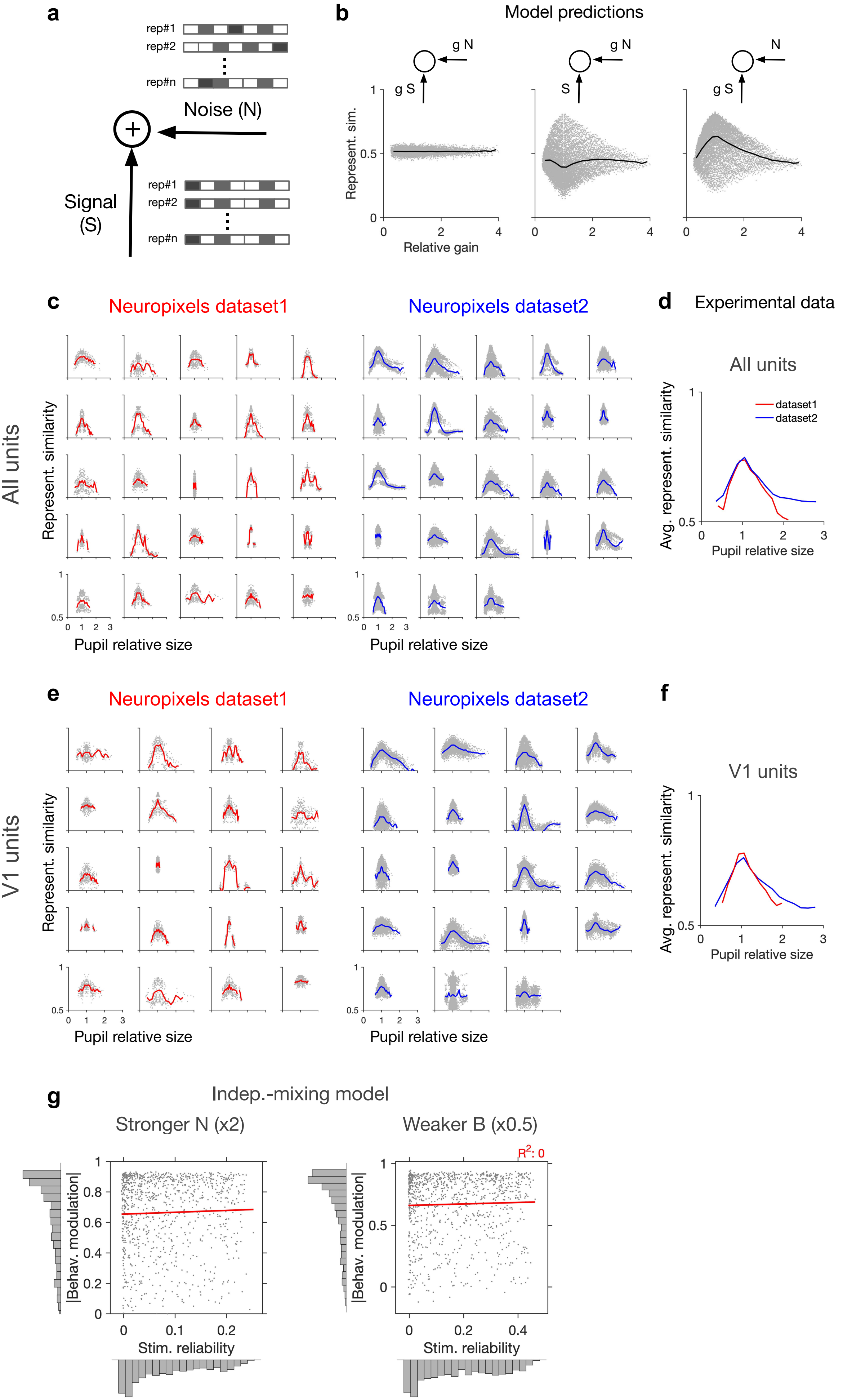

### Figure 2--figure supplement 2

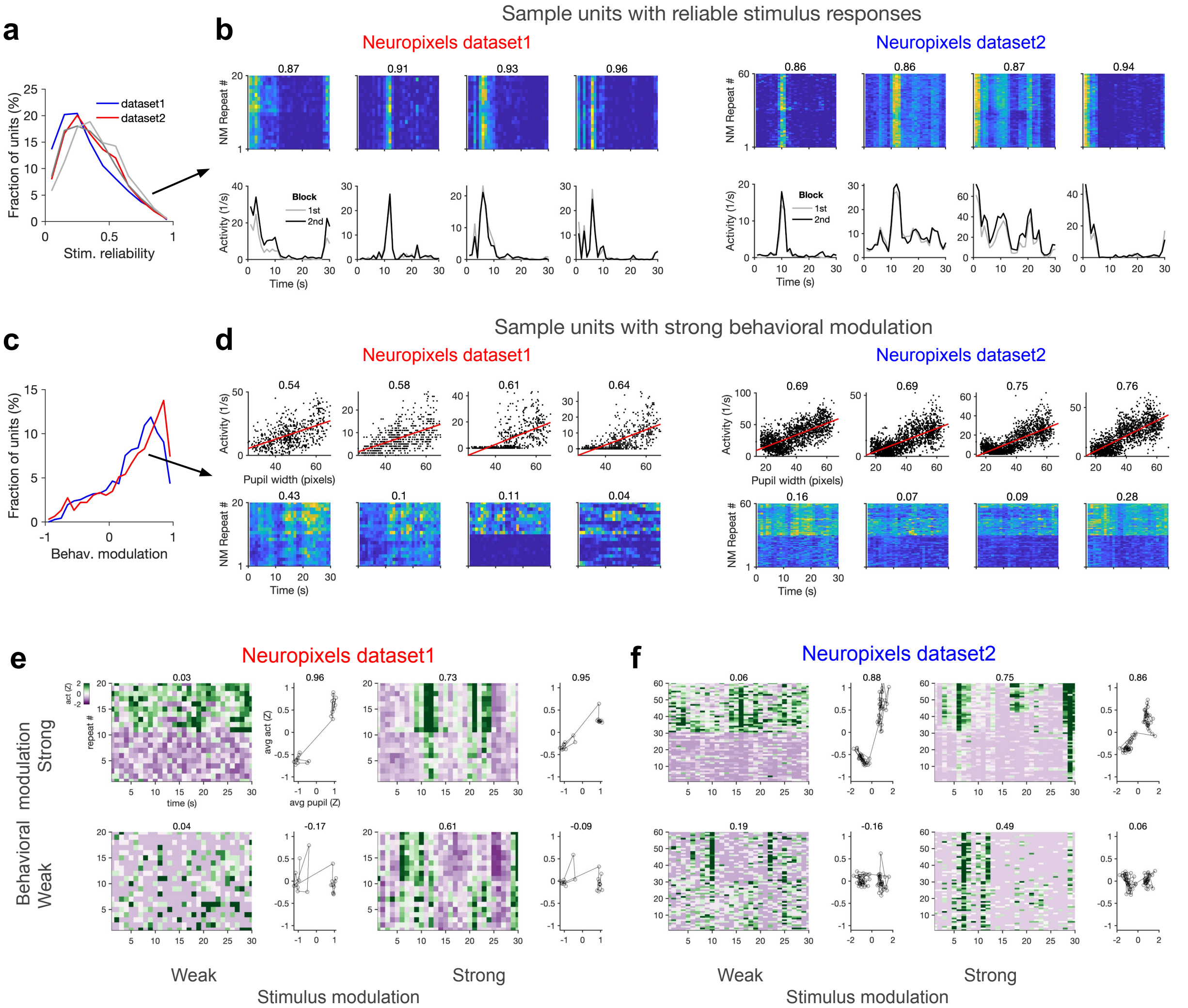

### Figure 2--figure supplement 3

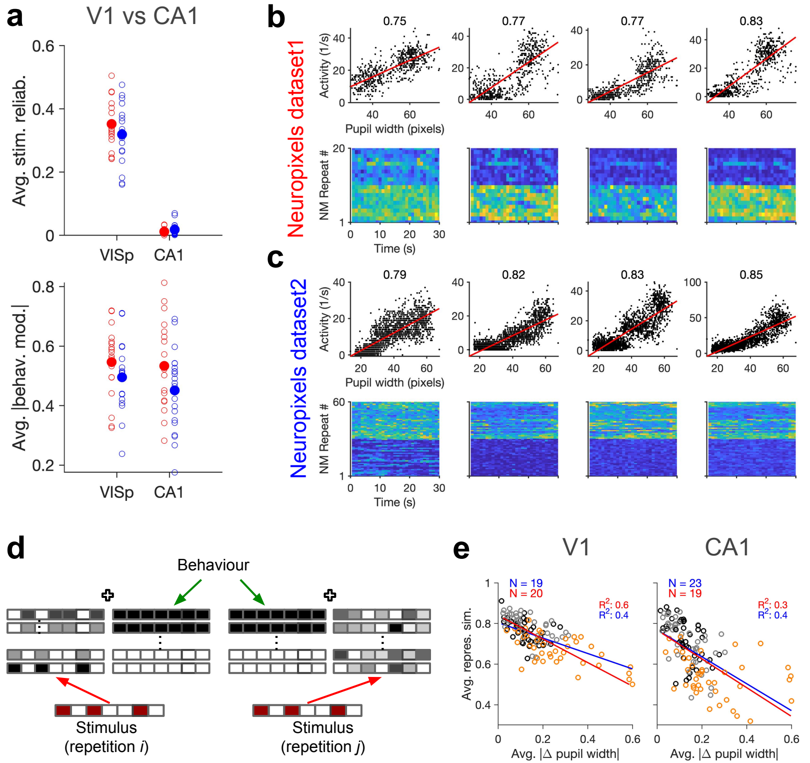

### Figure 3--figure supplement 1

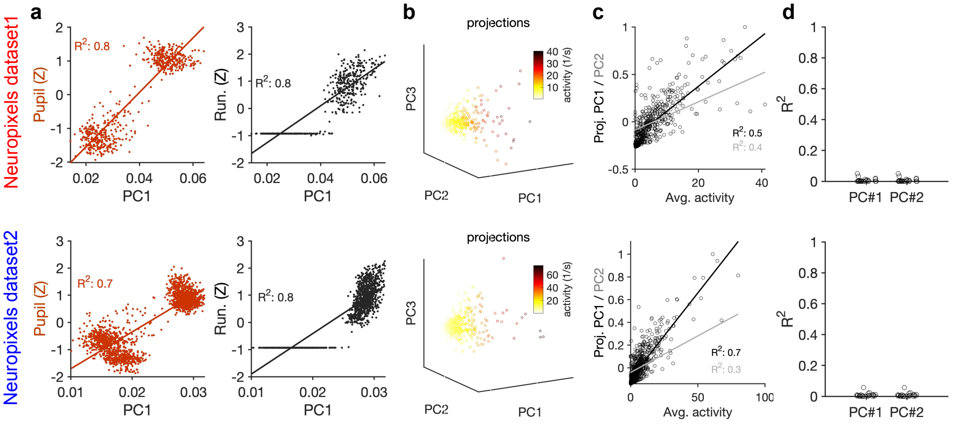

### Figure 3--figure supplement 2

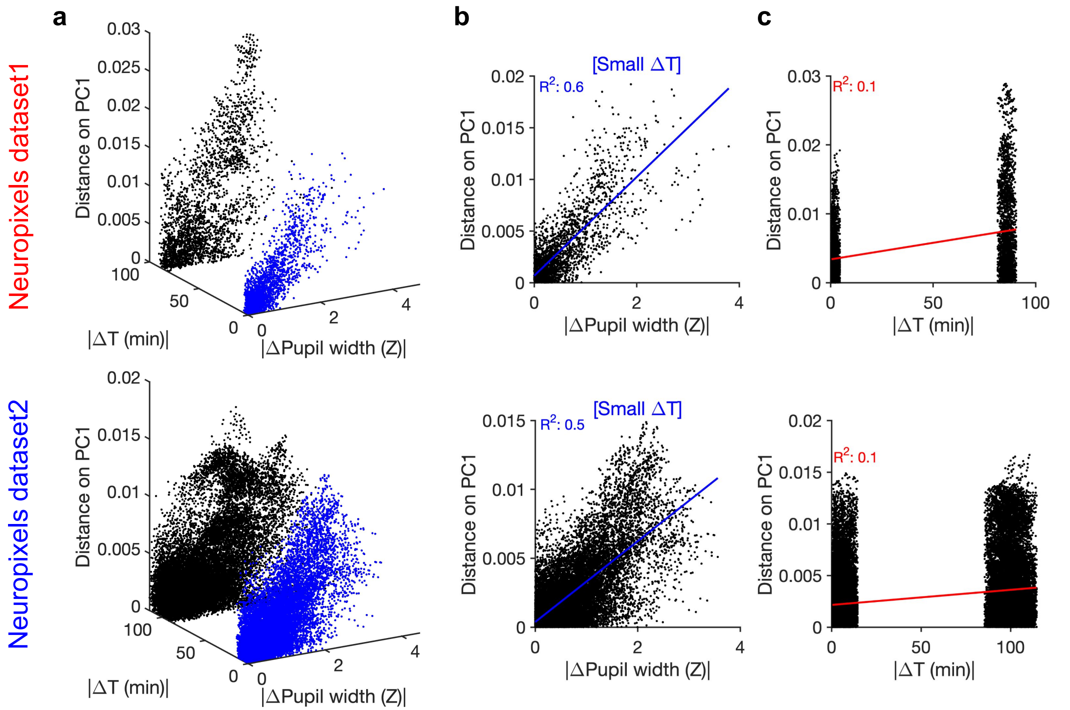

### Figure 4--figure supplement 1

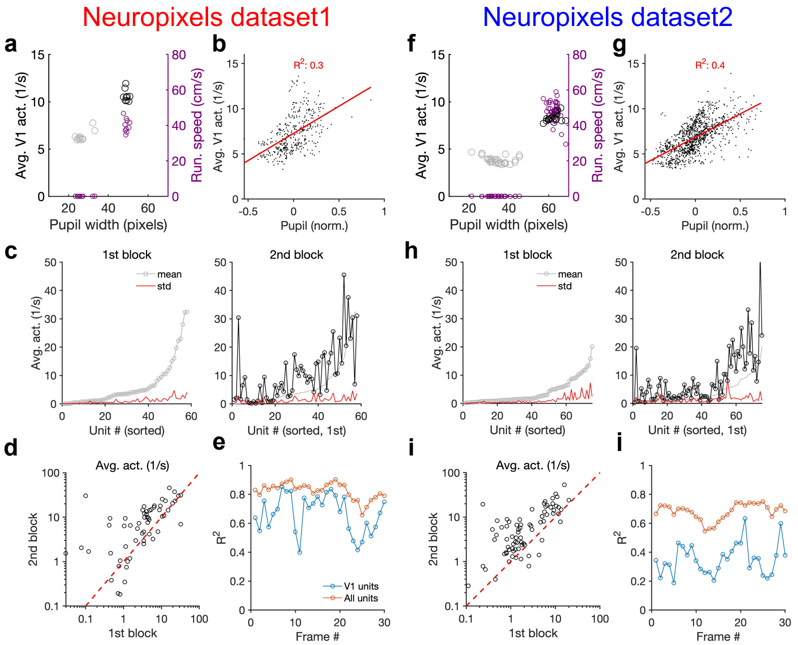

### Figure 4--figure supplement 2

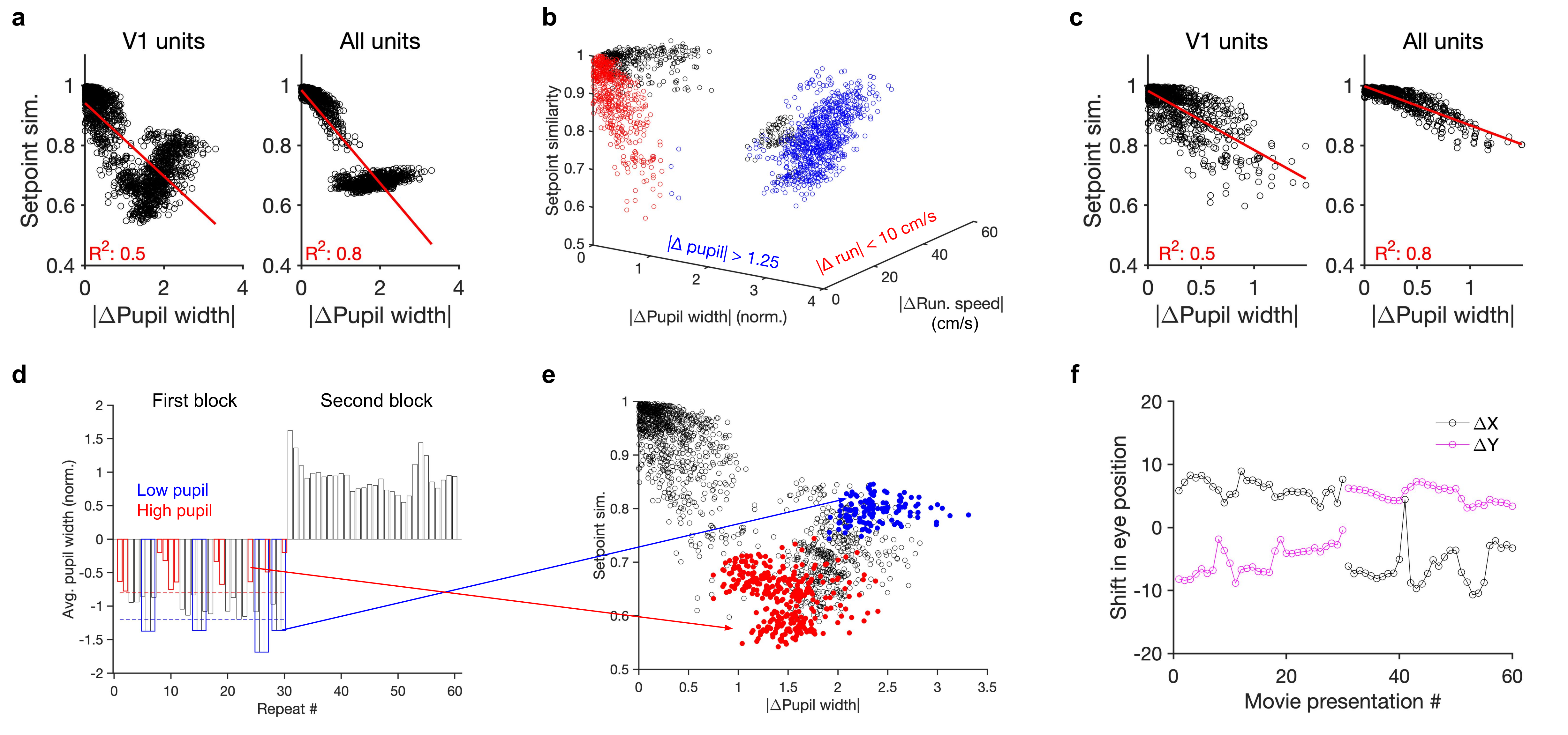

### Figure 4--figure supplement 3

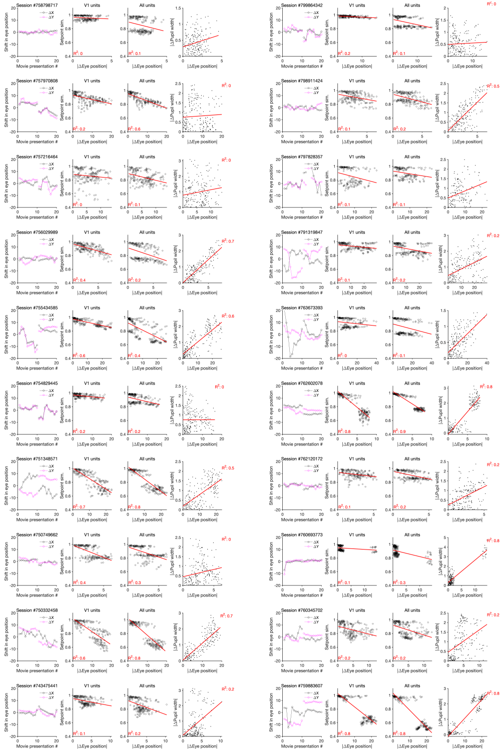

### Figure 4--figure supplement 4

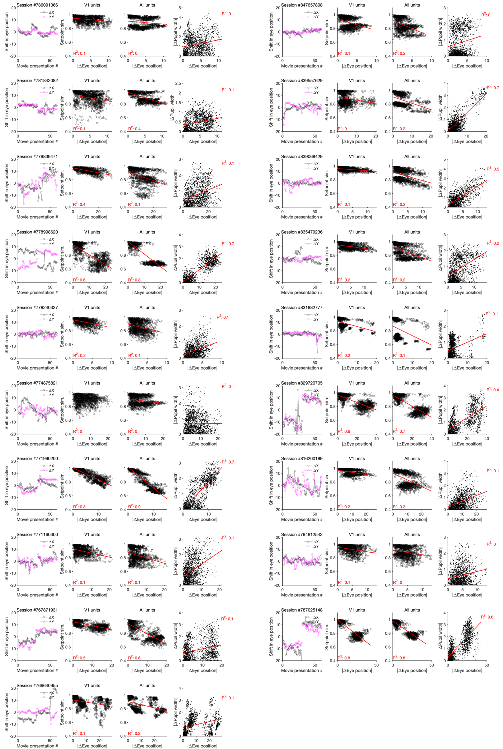

### Figure 4--figure supplement 5

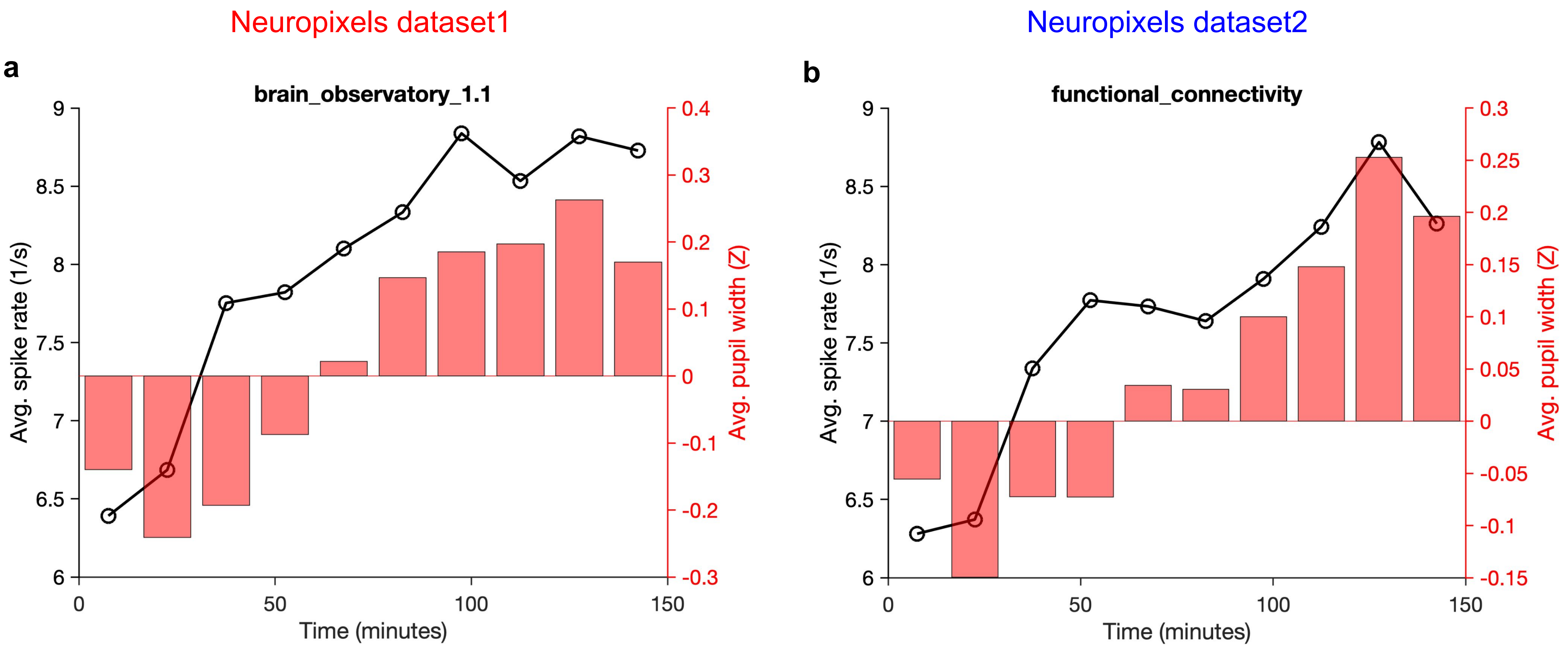

### Figure 5--figure supplement 1

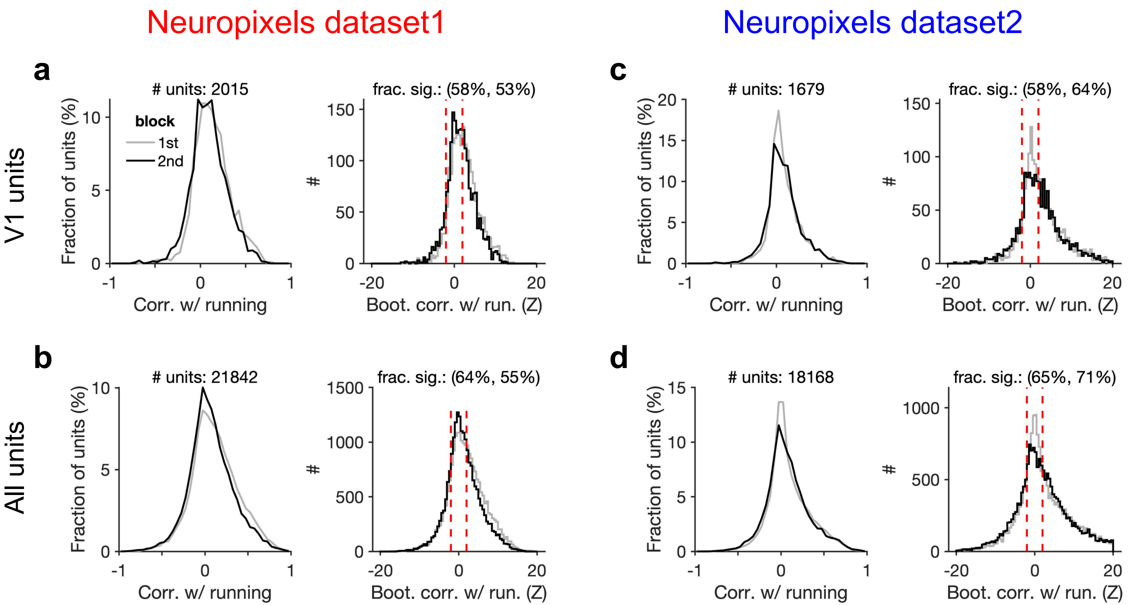

### Figure 5--figure supplement 2

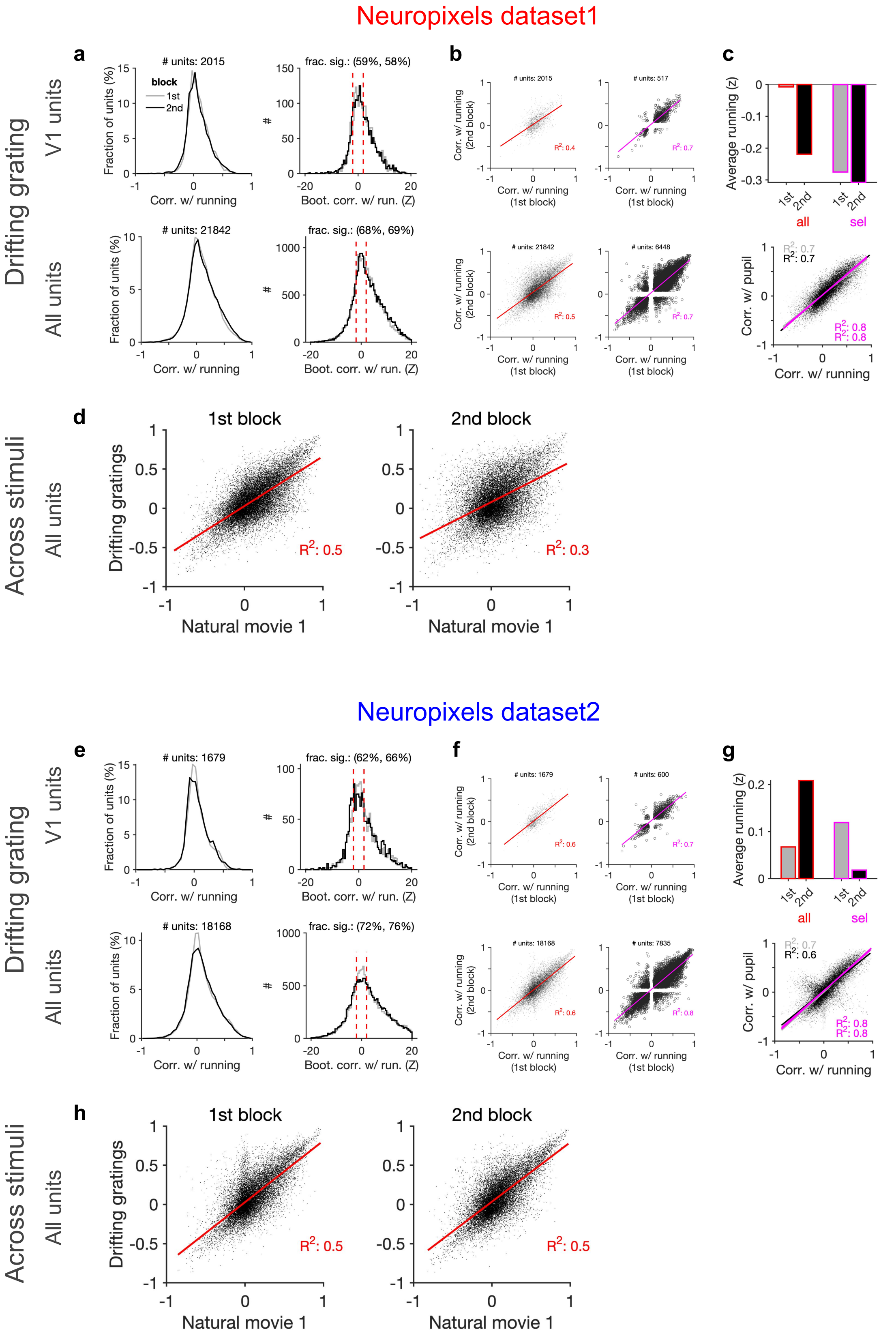

### Figure 5--figure supplement 3

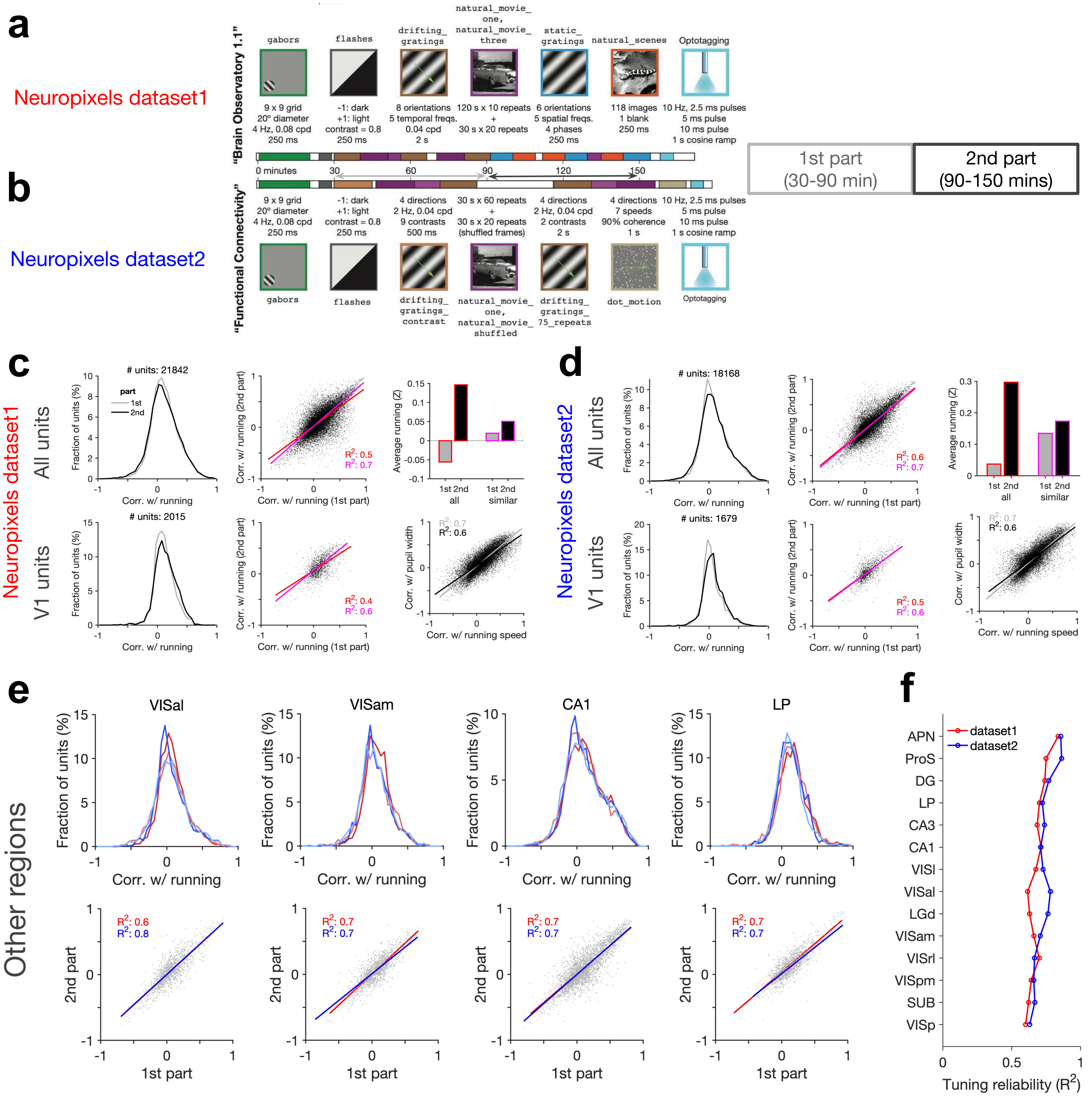

### Figure 7--figure supplement 1

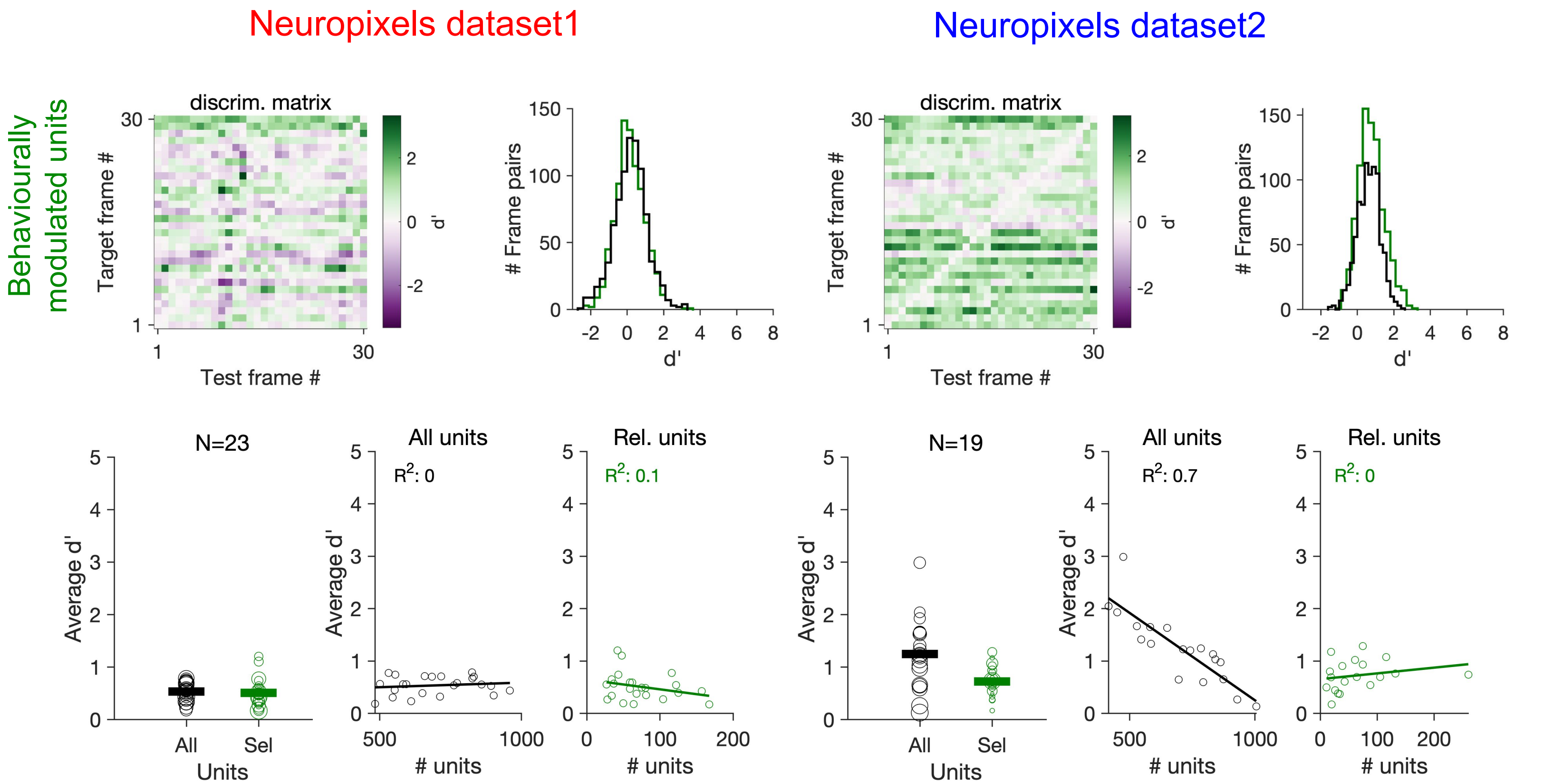

### Figure 7--figure supplement 2

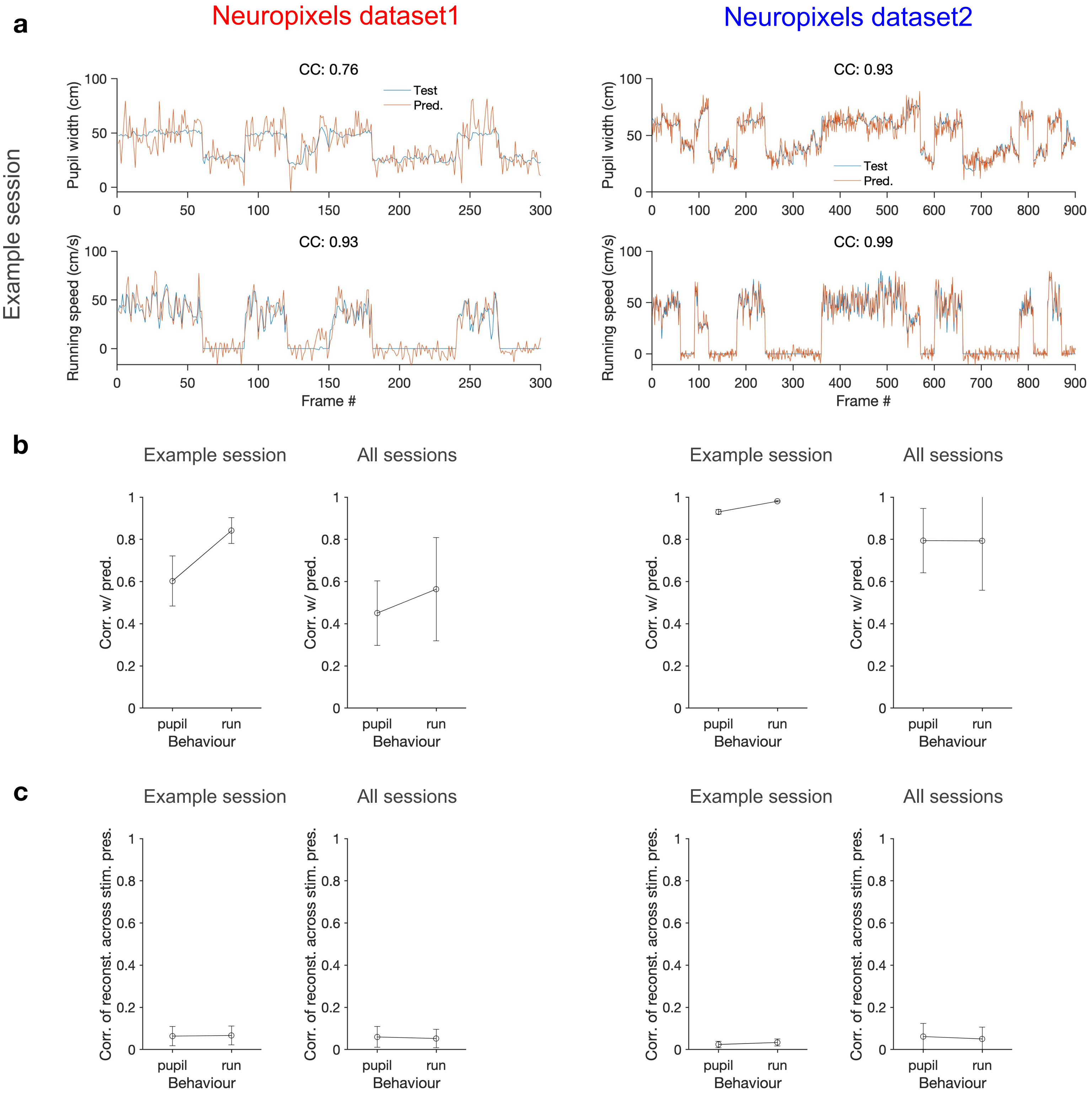
